# Enteric bacterial infection in *Drosophila* induces whole-body alterations in metabolic gene expression independently of the Immune Deficiency (Imd) signalling pathway

**DOI:** 10.1101/2022.05.11.491537

**Authors:** Rujuta Deshpande, Byoungchun Lee, Savraj S Grewal

## Abstract

When infected by intestinal pathogenic bacteria, animals initiate both local and systemic defence responses. These responses are required to reduce pathogen burden and also to alter host physiology and behaviour to promote infection tolerance, and they are often mediated through alterations in host gene expression. Here, we have used transcriptome profiling to examine gene expression changes induced by enteric infection with the gram-negative bacteria *Pseudomonas entomophila (P*.*e)* in adult female *Drosophila*. We find that infection induces a strong upregulation of metabolic gene expression, including gut and fat body-enriched genes involved in lipid transport, lipolysis, and beta-oxidation, as well as glucose and amino acid metabolism genes. Furthermore, we find that the classic innate immune deficiency (Imd)/Relish/NF-KappaB pathway is not required for, and in some cases limits, these infection-mediated increases in metabolic gene expression. We also see that enteric infection with *P*.*e*. down regulates the expression of many transcription factors and cell-cell signaling molecules, particularly those previously shown to be involved in gut-to-brain and neuronal signaling. Moreover, as with the metabolic genes, these changes occurred largely independent of the Imd pathway. Together, our study identifies many metabolic, signaling and transcription factor gene expression changes that may contribute to organismal physiological and behavioural responses to enteric pathogen infection.

## Introduction

When infected with pathogenic bacteria, animals need to mount appropriate defence responses to promote their survival. Perhaps the best-studied mechanisms are the innate immune responses that are involved in sensing invading pathogens and initiating antibacterial responses (Buchon *et al*. 2014). These mechanisms are termed resistance responses and are initiated to reduce pathogen burden. Recent work has also shown how bacterial infections can induce systemic changes in physiology and metabolism (Medzhitov *et al*. 2012; Martins *et al*. 2019; Troha and Ayres 2020). These metabolic changes can be required to fuel the energetically costly innate immune resistance responses to bacterial infection (Ryan and O’Neill 2020). In some cases, however, these changes can promote infection survival without reducing pathogen levels. These are termed tolerance responses and often involve changes in host metabolism, physiology, and behaviour (Ayres and Schneider 2012; Medzhitov *et al*. 2012) that function to limit the damaging effects of the pathogen and maintain host health. While these tolerance responses are not as well studied as the classic innate immune resistance responses, it is becoming clear that they are as important in determining individual susceptibility to infection (Ayres 2020).

*Drosophila* has been a versatile and informative model system in the investigation of organismal defence responses to bacterial infection (Younes *et al*. 2020). Oral infection of adult flies with gram-negative pathogenic bacteria induces both local and systemic resistance responses. Central to these responses is induction of the Imd/NF-Kappa B signalling pathway (Kleino and Silverman 2014). This pathway senses bacteria through cell surface peptidoglycan recognition protein (PGRP) receptors and signals through an intracellular cell signalling pathway involving the death domain containing protein, Imd, leading to transcriptional activation of the Relish/NF Kappa B transcription factor (Capo *et al*. 2016). One main target is of Relish are antimicrobial peptides that mediate resistance responses to reduce pathogen load (Buchon *et al*. 2014). Enteric gram-negative bacterial infection in *Drosophila* can also induce both local and systemic changes in host physiology and metabolism to promote both resistance and tolerance response (Lee and Lee 2018; Galenza and Foley 2019; Bland 2022). For example, gut infection induces altered lipid storage and metabolism both in the intestine and in remote tissues such as fat body and oenocytes (Hang *et al*. 2014; Charroux *et al*. 2018; Kamareddine *et al*. 2018a; Kamareddine *et al*. 2018b; Lee *et al*. 2018; Harsh *et al*. 2019; Zhao and Karpac 2021; Charroux and Royet 2022; Deshpande *et al*. 2022). Enteric infection can also induce alterations in systemic carbohydrate, mitochondrial and amino acid metabolism (Zhao and Karpac 2021). These non-autonomous effects of enteric infection on whole-body metabolism involve changes in both gut-to-fat and muscle-to-fat metabolic signalling and are required to promote host fitness. Enteric bacterial infection can also cause changes in fly behaviour that may help to promote infection tolerance, such as food avoidance and reduced fecundity (Soldano *et al*. 2016; Masuzzo *et al*. 2019; Charroux *et al*. 2020; Masuzzo *et al*. 2020). In some cases, these effects have been shown to occur as a result of altered gut-to-brain signalling (Cai *et al*. 2021). Together, these studies show that gut pathogens can trigger tissue-to-tissue signalling in flies to coordinate changes in metabolism, physiology and behaviour and promote both resistance and tolerance responses to infection. Understanding the mechanisms underlying these changes will provide further insights into how organismal defence responses.

One way that organismal physiology can be altered is through changes in gene expression. To provide further insight into systemic responses to enteric infection, we have used transcriptome profiling to identify whole-body gene expression changes following enteric infection with *P*.*e*. Our data reveal upregulation of many metabolic genes involved in lipid, carbohydrate, and amino acid metabolism, and we see that induction of these genes occurs independently of Imd signaling and, in some cases, is antagonized by the Imd signaling pathway. In addition, we saw a downregulation of genes encoding transcription factors, signalling peptides, and signalling receptors, particularly those involved in gut-to-brain and neuronal signaling. Together, these analyses reveal broad changes in metabolic, signaling and transcription factor gene expression that may mediate organismal responses to infection.

## Results and Discussion

### Enteric infection upregulates the expression of metabolic genes

Adult mated females were either mock-infected (fed with sucrose alone) or orally infected by feeding with *P*.*e*. We then isolated whole-body RNA at 16 hours post-infection for RNA seq analysis (Figure 1A). Using a cut-off of +/-1.5 fold and a false-discovery rate of p<0.05, we identified 1233 transcripts showing increased expression following infection and 1602 transcripts showing reduced expression (Figure 1B). The best-studied immune response to gram negative infection in *Drosophila* involves upregulation of the Imd/NF-Kappa B signalling pathway. Among the most strongly upregulated genes were the antimicrobial peptides (AMPs) which are known targets of the Imd pathway, and which function to reduce pathogen burden and promote resistance responses (Figure 1C). We also saw induction of several other classes of genes encoding for secreted peptides and proteases that have also previously been shown to be induced upon bacterial infection and to mediate anti-microbial responses (Figure 1C). These results confirm that our infection protocol induces a robust transcriptional immune response to enteric infection.

**Figure 1.**
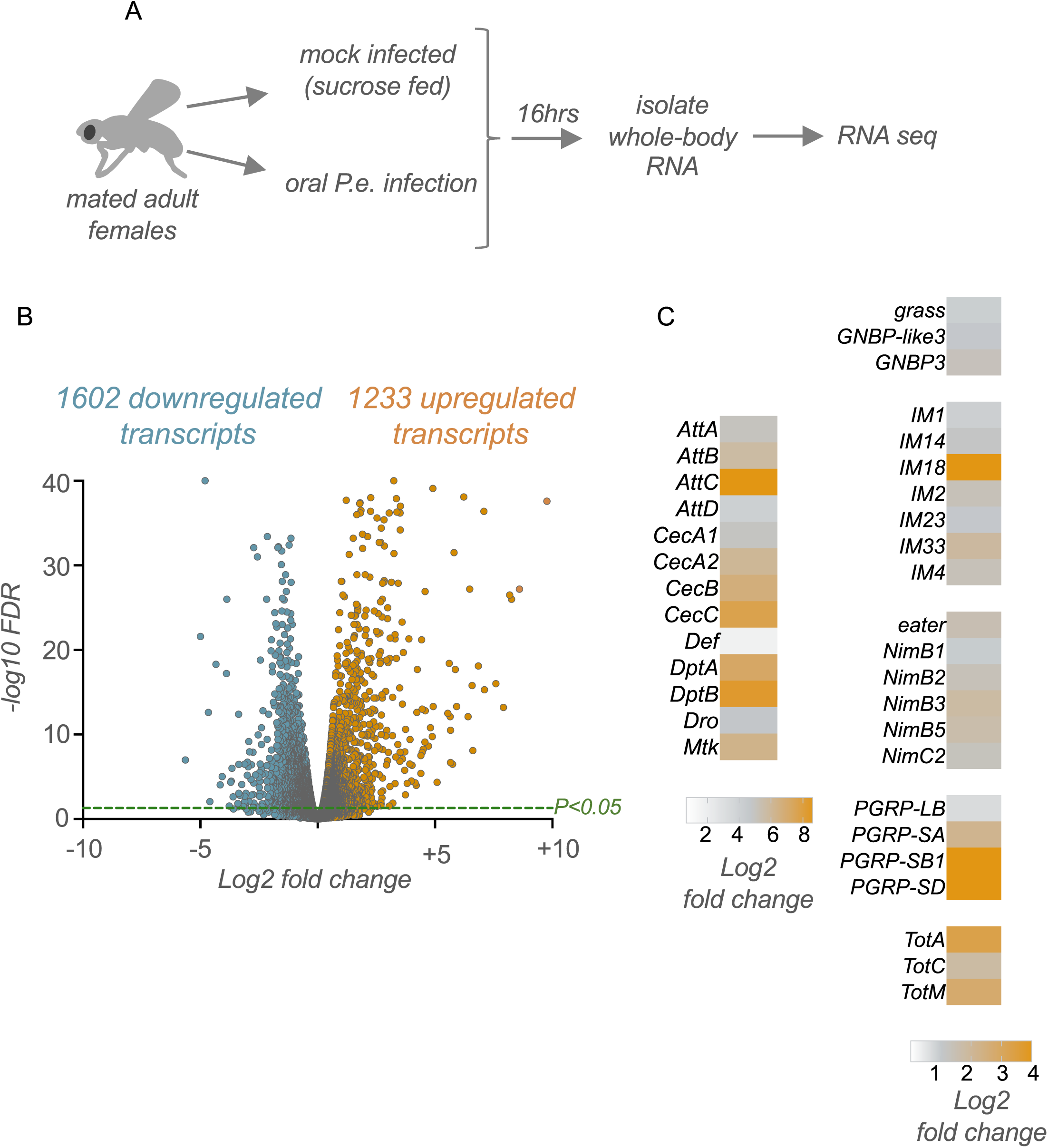
Enteric *P*.*e*. infection induces alterations in whole-body gene expression including upregulation of antimicrobial peptide genes. A) Schematic outline of our experimental approach. B) Volcano plot showing the up-and down-regulated genes following enteric infection. C) Heatmap depicting the change in expression (Log2 fold change vs mock-infected flies) of antimicrobial peptide (AMP) genes and other immune-response genes following enteric infection.

We then carried out KEGG and Gene Ontology analyses of the upregulated transcripts to help identify functional classes of genes whose expression is induced upon infection. As anticipated, KEGG pathway analyses showed enrichment in Toll and Imd signalling, while GO analysis showed the most significant enrichment was in the biological process categories of defense response and immune system process genes (figure 2A, B). The other main classes of upregulated genes were, interestingly, related to metabolism. These included genes involved in lipid, carbohydrate, and amino acid metabolism (Figure 2B, C). Previous transcriptomic studies following enteric infection also showed changes in metabolic genes in specific tissues such as intestine, thoracic muscle, or abdominal adipose tissues (Buchon *et al*. 2009; Zhao and Karpac 2021). Similarly, transcriptome studies using models of systemic infection in flies also showed changes in metabolic gene expression (Clark *et al*. 2013; Troha *et al*. 2018). Our work adds to these studies to suggest that remodelling of host metabolism through altered gene expression is a common response to pathogenic bacterial infection in flies.

**Figure 2.**
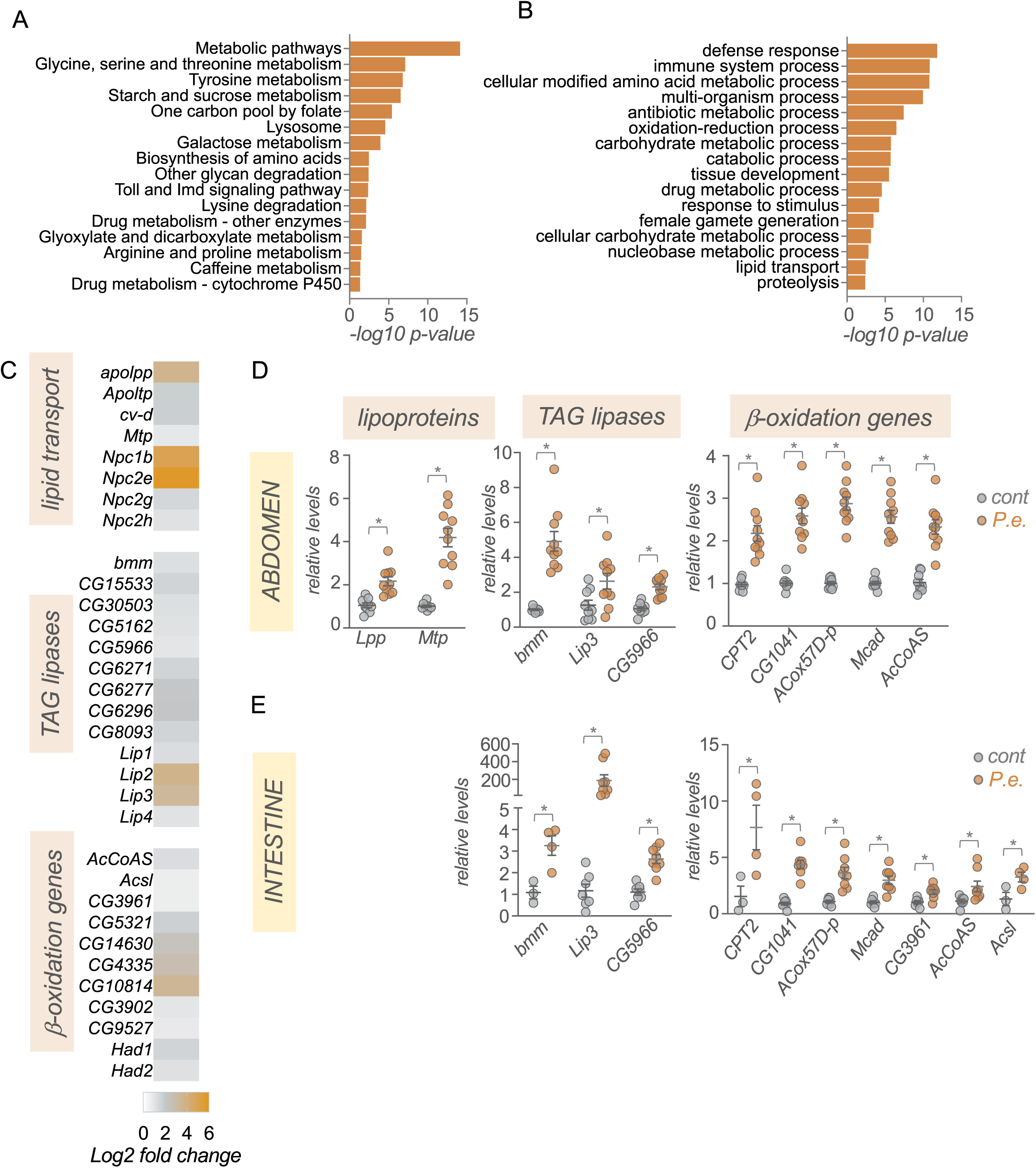
Enteric *P*.*e*. infection leads to upregulation of gut-and fat body-expressed lipid transport, TAG lipases and beta-oxidation genes. A) KEGG pathway analysis of genes showing >1.5-fold increase following enteric infections. B) GO analysis (biological process categories) of genes showing >1.5-fold increase following enteric infections. C) Heatmap depicting the change in expression (Log2 fold change vs mock-infected flies) of lipid transport, TAG lipases and beta-oxidation genes following enteric infection. Grey symbols show genes with strong enrichment in either the intestinal tissues or fat body, based on expression levels from FlyAtlas2. D, E) qPCR analysis of lipid transport, TAG lipases and beta-oxidation genes from, D, isolated abdominal tissues or E, intestines. Bars represent mean +/-SEM. Symbols represent individual data points, n=6-10 per condition. * p<0.05, Students t-test.

We examined the changes in metabolic gene expression by first exploring lipid metabolism genes. We saw increased expression of genes involved in lipid transport such as the Niemann-Pick type C family of cholesterol transporters, and the apolipoproteins, *Lpp, Ltp, cv-d* and *Mtp* that are used to transport lipids from lipid storage tissues, such as the gut and abdominal adipose tissues, to other organs (Palm *et al*. 2012)(Figure 2C). We also saw increased whole-body expression of TAG lipases and beta-oxidation genes (Figure 2C). This is consistent with reports showing the depletion of lipid stores following enteric infection (Hang *et al*. 2014; Kamareddine *et al*. 2018a; Kamareddine *et al*. 2018b; Zhao and Karpac 2021; Deshpande *et al*. 2022). The main lipid storage tissues in the adult fly are the intestine and the fat body and oenocytes which are enriched in the abdomen. We used qPCR to analyze gene expression in samples of isolated intestine and abdomen (abdominal tissues with intestine, ovaries and Malpighian tubes removed) from mock-infected vs *P*.*e*.-infected flies. Consistent with our RNAseq analyses, we saw increased expression of the lipoproteins *Lpp* and *Mtp* in the abdominal samples (Figure 2D). We also saw increased expression of both TAG lipases and beta-oxidation genes in both the intestine and abdominal samples (Figure 2D, E). Together these changes in gene expression suggest that upon enteric infection, flies alter their metabolism to mobilize fat and gut lipid stores and transport these lipids to other tissues to fuel metabolism through beta-oxidation. In fact, a recent report has described how differences in fat mobilization can explain inter-individual differences in susceptibility to infection (Zhao and Karpac 2021). The gut and fat body are key tissues that coordinate host immune response to infection. They do so in large part by both increasing expression and release of the AMPs, and by initiating organ-to-organ signalling to coordinate whole-body host defences responses (Buchon *et al*. 2009). These effects rely on increased endocrine functions of both tissues which imposes a high protein synthetic and secretory burden (Martinez *et al*. 2020). As a result, both the gut and fat body likely have high metabolic demands following infections. Our RNA seq and qRT-PCR data suggest that both tissues may upregulate fatty acid oxidation to sustain and support their metabolic needs. This is in line with emerging work in immunometabolism showing that many key immune system cells and tissues switch to fatty acid oxidation to meet their metabolic needs and to allow them to function properly to mediate defence responses (O’Neill *et al*. 2016; Cumnock *et al*. 2018; Batista-Gonzalez *et al*. 2019).

In addition to alterations in lipid mobilization and lipid metabolic genes, we also saw that enteric infection led to an upregulation of whole-body expression of genes encoding regulators of both carbohydrate and amino acid metabolism (Figure 3A-C). For example, we saw increased expression of genes involved in both glycogen mobilization, and trehalose synthesis and transport (*GlyP, UGP, Tps1, Tret1-1, Tret1-2*) as well as increased expression of gluconeogenesis/glycolysis (*tobi, Pepck1, Hex-C*) and pentose phosphate pathway genes (*Pgd, Zw, Taldo)* (Figure 3A). These changes suggest that flies remodel their glucose metabolism to adapt to infection. The upregulation of glycogen mobilization genes is consistent with previous reports showing depletion of glycogen stores following enteric infection (Hang *et al*. 2014; Deshpande *et al*. 2022). The muscle and fat body are the main glycogen storage tissues in flies, and one possibility is that the mobilized glucose may be used to fuel glycolysis and/or the pentose phosphate pathway in these tissues to help support their metabolic needs. In addition, the upregulation of trehalose synthesis genes suggests that some of the glucose may also be converted into trehalose, the main circulating form of glucose in flies, to be transported to other tissues.

**Figure 3.**
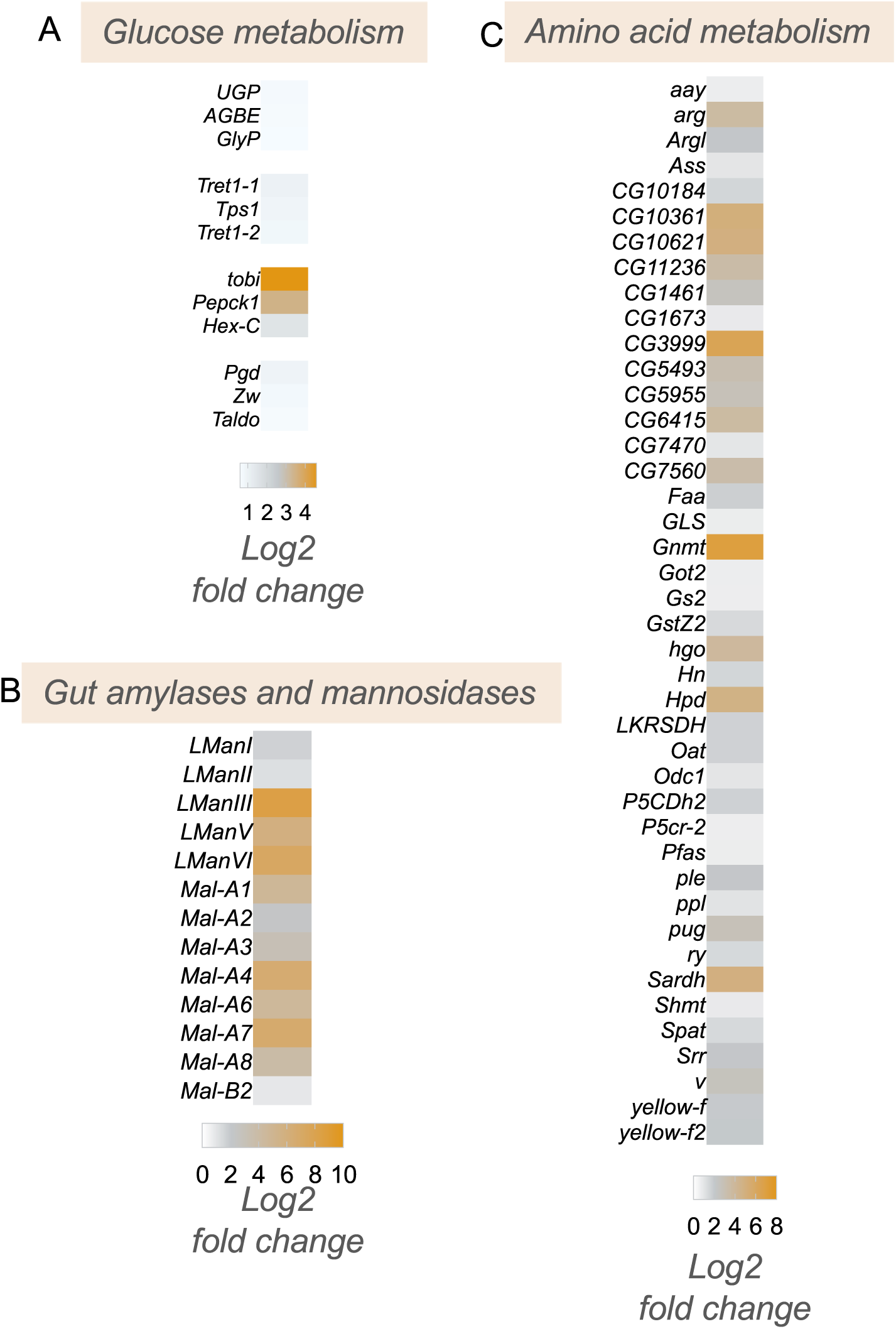
Enteric *P*.*e*. infection induces increase in sugar and amino acid metabolism genes. Heatmap depicting the change in expression (Log2 fold change vs mock-infected flies) of A, glucose metabolism, B, amylases and mannosidases, and C, amino acid metabolism genes following enteric infection.

Among the upregulated amino acid metabolism genes, two of the most strongly induced were *Gnmt* and *Sardh* (Figure 3C). Both are involved in regulation of methionine metabolism and the control of S-adenosylmethione (SAM) levels. SAM is an important metabolite since it functions as a universal methyl donor in the control of methyltransferase activity. Interestingly, previous studies have shown that regulation of SAM levels in both the intestine and fat body play important roles in regulating physiology and tissue homeostasis. For example, alterations in SAM levels in the intestine have been shown to regulate levels of the Upd3 cytokine to control stem-cell mediated tissue renewal (Obata *et al*. 2018), which is known to be an important tissue repair response following enteric pathogen infection (Buchon *et al*. 2009; Jiang *et al*. 2009). In addition, alterations in SAM levels in the fat body have been shown to occur following necrotic wing injury and this alteration is important for controlling lipid levels and survival (Obata *et al*. 2014). Increased fat body expression of Gnmt and modulation of SAM levels can also extend lifespan (Obata and Miura 2015; Tain *et al*. 2020). Based on these previous reports, our findings suggest that modulation of intestine and/or fat body SAM levels may be an important metabolic response to enteric bacterial infection.

### The Imd pathway antagonizes the infection-mediated upregulation of metabolic gene expression

We next explored potential signalling and transcriptional mechanisms that might explain the upregulation in metabolic gene expression following enteric pathogen infection. We focused particularly on examining the conserved Imd/NF-Kappa B innate immunity pathway. The Imd pathway is the main signalling pathway induced by pathogenic gram-negative infection in *Drosophila* (Kleino and Silverman 2014). The pathway is activated when cell-surface PGRP receptors detect bacterial peptidoglycans and stimulate a downstream intracellular signalling cascade involving Imd, a death-domain containing protein, that eventually leads to the activation and nuclear localization of the NF-Kappa B transcription factor, Relish (Kleino and Silverman 2014). One of the main transcriptional targets of Relish are the AMPs, which mediate the main antibacterial immune response to reduce pathogen load. We used qRT-PCR to compare whole-animal gene expression levels in control (*w*^*1118*^) or *imd* mutant animals that had either been mock-infected or orally infected by 24 hr feeding with *P*.*e*. We saw that the infection-mediated upregulation of two AMPS, *CecA* and *CecC* seen in control (*w*^*1118*^) animals was completely suppressed in the *imd* mutant animals, confirming that this line is a strong loss-of-function of the Imd pathway (Figure 4A). We then analysed expression of representative metabolic genes from several main classes: lipoproteins, TAG lipases, beta oxidation genes, carbohydrate metabolic genes, and amino acid metabolic genes. From these analyses, three main findings emerged (Figure 4B-F). First, we found that infection induced a strong upregulation of all the metabolic genes that we tested, thus confirming the transcript changes that we saw in our RNA seq analyses. Second, we saw that none of these increases in metabolic gene expression were blocked in the *imd* mutants. Thus, in contrast to the induction of the AMPs, the increase in metabolic gene expression does not rely on the Imd/NF-Kappa B immune signalling pathway. Thirdly, and most interestingly, we saw that in many cases, the increase in metabolic gene expression seen following infection was further exacerbated in the *imd* mutant flies. These results suggest that the Imd pathway may function to antagonize the infection-mediated changes in host metabolism. Indeed, a previous report showed that constitutive genetic activation of the Imd pathway in larval fat body led to downregulation of many carbohydrate and lipid metabolic genes, including several that we see are induced upon infection and further increased in infected *imd* mutants (Davoodi *et al*. 2019). In addition, Relish, the transcriptional effector of the Imd pathway, has been shown to antagonize the starvation-mediated induction of the lipase, *bmm* (Molaei et al. 2019). Moreover, Relish knockdown has been shown to exacerbate the infection-induced increase in amino acid metabolism genes in muscle, including Gnmt, and the extent of these muscle effects of Relish were shown to underlie inter-individual differences in infection susceptibility (Zhao and Karpac 2021). These previous reports together with our RNA seq and qPCR results suggest that while induction of the Imd pathway is necessary to induce AMPs and induce anti-bacterial responses, it also functions to limit induction of metabolic genes.

**Figure 4.**
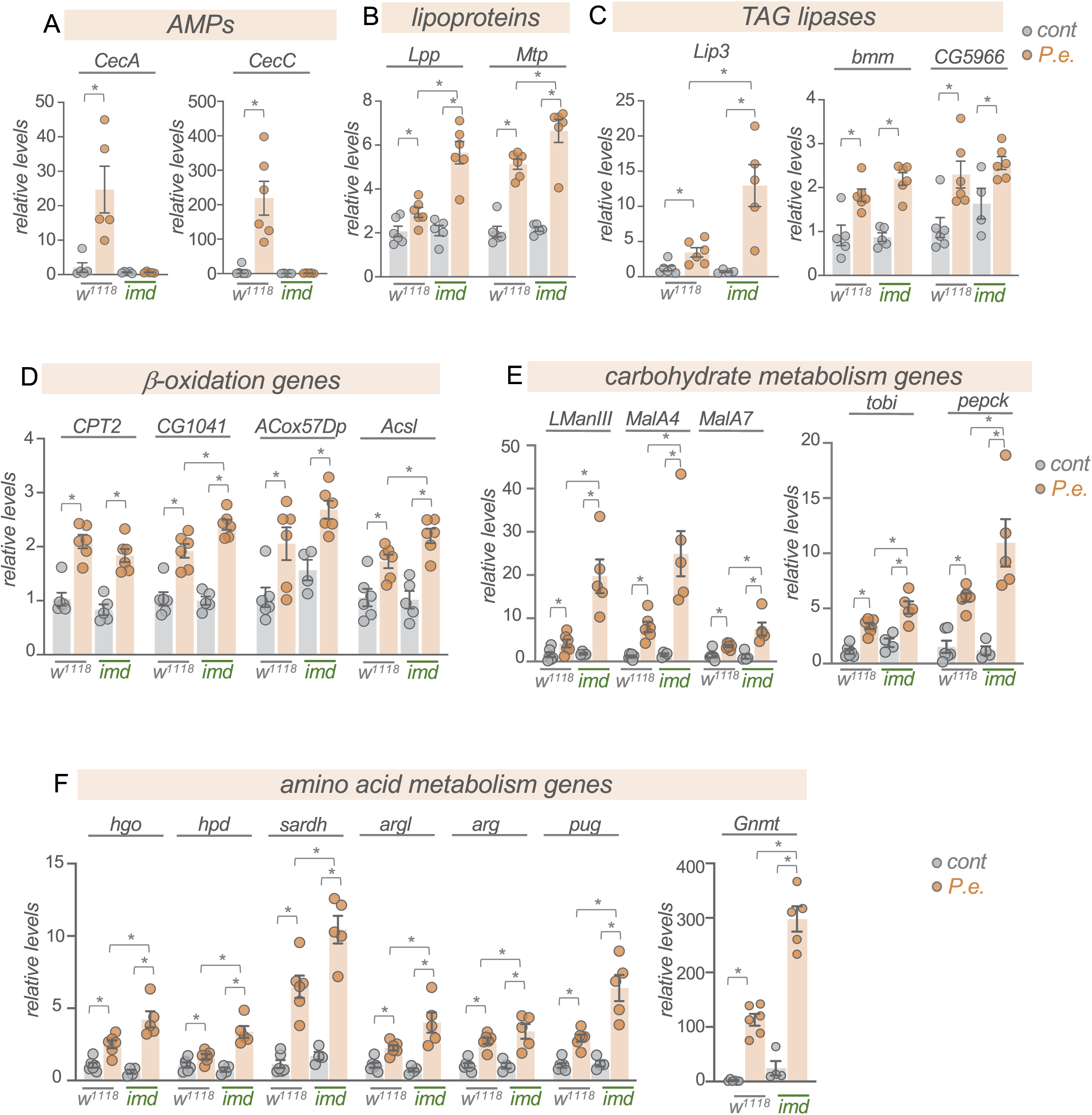
Enteric *P*.*e*. infection induces expression of metabolic genes independently of the Imd pathway. qRT-PCR analysis of mRNA levels of A) AMPs, B) lipoprotein genes, C) TAG lipases, C) beta-oxidation genes, E) carbohydrate metabolism genes, and F), amino acid metabolism genes from control (*w*^*1118*^) vs *imd* mutant adult females, that were either mock-infected (grey bars and symbols) or infected by oral feeding with *P*.*e*. for 24 hours (orange bars and symbols). Bars represent mean +/-SEM. Symbols represent individual data points, n=4-6 per condition. * p<0.05, Two-way ANOVA, followed by Students t-test.

What signalling pathways and transcriptional mechanisms, if not the Imd pathway, may lead to the upregulation of metabolic gene expression following infection? One possibility is the endocrine insulin/PI3K/TOR pathway. We previously showed that this pathway is induced upon enteric infection with *P*.*e*. and we found that this induction was needed to promote lipid synthesis gene expression (Deshpande *et al*. 2022). Similarly, Charroux and Royet showed that infection induces an upregulation of SREBP in the fly fat body through insulin/PI3 kinase signalling (Charroux and Royet 2022). Thus, given that the insulin/P3K/TOR pathway is the one of the main regulators of organismal metabolism in flies, many of the increases in lipid, carbohydrate, and amino acid metabolism genes that we see maybe mediated through this pathway. Furthermore, since Imd signalling has been shown to negatively regulate insulin/PI3K signalling (Davoodi *et al*. 2019), this may also explain, in part, why the increased expression of metabolic genes is further exacerbated in *imd* mutants. It may seem paradoxical that insulin/PI3K/TOR signalling would induce both lipid synthesis and lipolysis genes. However, it is possible that these effects may be occurring in a cell-or tissue-specific manner. For example, activation of the PI3K/TOR pathway in oenocytes has been shown to reduce lipid levels in these cells, but then lead to an increase in lipid levels in fat body cells (Ghosh *et al*. 2020).

### Enteric infection downregulates the expression of many CNS and intestinal signalling pathways and transcription factors

We saw that 1602 transcripts showed reduced mRNA expression following infection (Figure 1B). GO term and KEGG pathway analyses of these genes, interestingly, revealed enrichment in two broad classes of genes – transcription factors and cell-cell signalling pathways (Figure 5A, B). We found that ∼100 transcription factors showed reduced expression following infection (Figure 5C). We used FlyAtlas2 (Leader *et al*. 2018) to explore where each of these transcription factors is normally expressed, and we saw that many of them show enriched expression either in the intestinal system (crop, midgut and/or hindgut) or the head and brain (head, eye, brain/CNS, and/or thoracicoabdominal ganglion)(Figure 5C). We also selected a few of these genes and examined whether their upregulated expression was mediated through the Imd pathway by using qRT-PCR to compare infection-mediated changes in whole-body expression in control (*w*^*1118*^) vs *imd* mutants. We saw that for each of the six transcription factors that we examined (*bap, Doc1, tll, grn, rib* and *sna)* mRNA expression levels were significantly reduced following infection, thus confirming the transcript changes that we saw in our RNA seq analyses. (Figure 5D). We also saw that expression levels of each transcription factor were significantly lower in mock-infected *imd* mutants compared to controls, and that infection did not decrease these levels further (except for *sna)* (Figure 5D). These results suggest that the infection-mediated decreases in expression of these transcription factors is not mediated through induction of Imd signaling.

**Figure 5.**
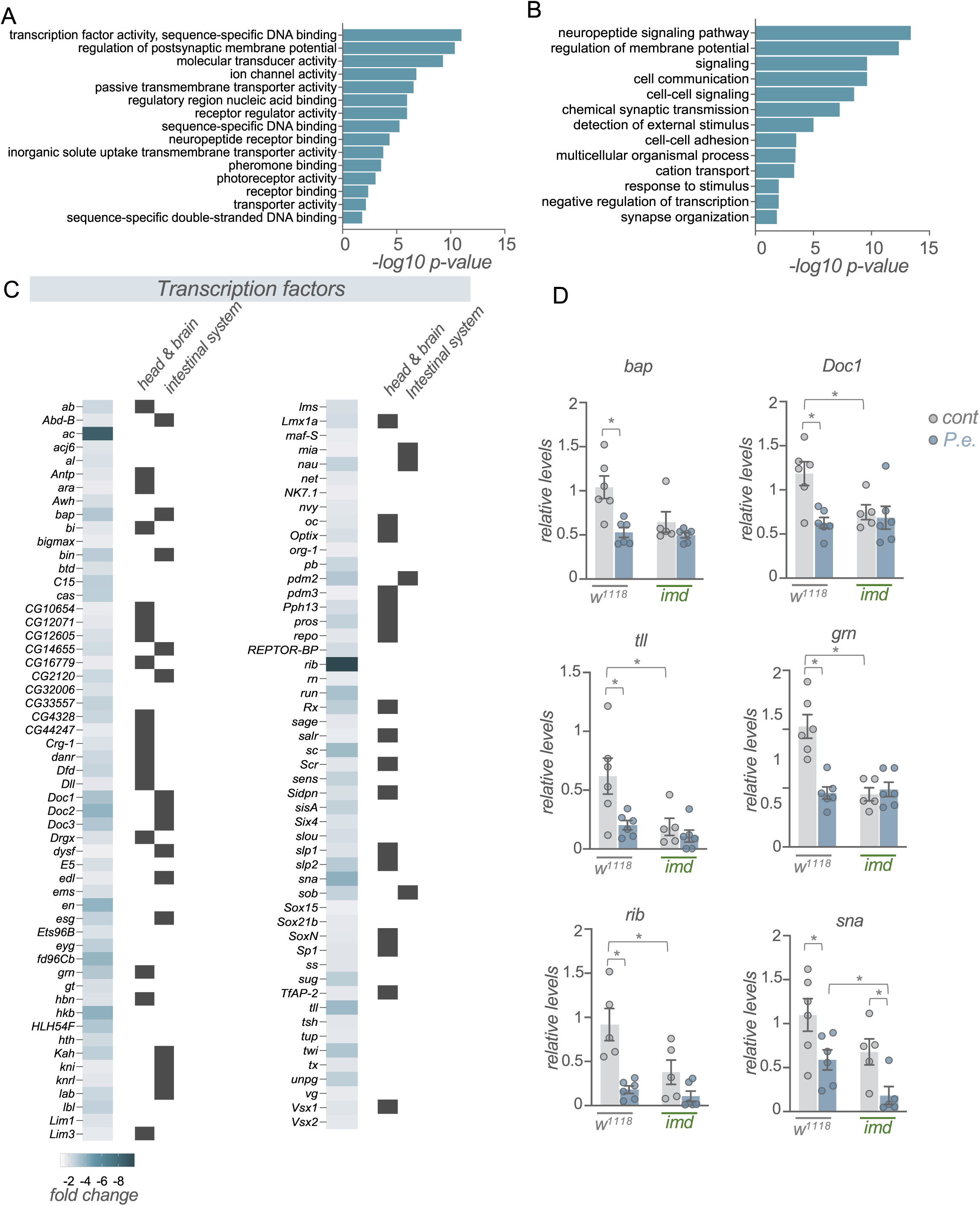
Enteric *P*.*e*. infection downregulates the expression of many transcription factor genes. A, B) GO analysis (A, biological process categories, B, molecular function categories) of genes showing >1.5-fold increase following enteric infections. C) Heatmap depicting the change in expression (Log2 fold change vs mock-infected flies) of transcription factor genes following enteric infection. Grey symbols show genes with strong enrichment in either the head/brain or intestinal system, based on expression levels from FlyAtlas2. D). qPCR analysis of selected transcription factor mRNA levels from control (*w*^*1118*^) vs *imd* mutant adult females, that were either mock-infected (grey bars and symbols) or infected by oral feeding with *P*.*e*. for 24 hours (blue bars and symbols). Bars represent mean +/-SEM. Symbols represent individual data points, n=4-6 per condition. * p<0.05, Two-way ANOVA, followed by Students t-test.

Another large group of genes that showed reduced expression following infection are those involved in cell-cell signaling (Figure 5A, B). These include both cell surface receptors (Figure 6A) and secreted peptide ligands (Figure 6B). In some cases (e.g., *AstA, Capa, FMRFa, Ms, Trissin, Tk)* we found expression of both the secreted factors and as well as their corresponding receptors were coordinately decreased, suggesting downregulation of signaling through pathways coupled to these ligand/receptor pairs. Also, as with the downregulated transcription factors, we found that most downregulated receptors and ligands showed enriched expression in the head/CNS and/or intestinal system as indicated by FlyAtlas2 expression profiles (Figure 6A, B). We selected several of these signaling peptides (*Gpa2, hug, pdf, proc, FMRFa, trissin, wg*) to examine whether the infection-mediated decrease in expression was dependent on the Imd signaling pathway. Using qRT-PCR analysis, we found that the whole-body expression of each of these genes was decreased following enteric infection with *P*.*e*., consistent with our RNA seq analyses (Figure 7). For three of these genes (*Gpa2, hug and pdf)* the expression levels in both the mock-infected and infected *imd* mutants were significantly lower than the control mock-infected flies, suggesting that the infection-mediated down-regulation of these genes did not occur through increased Imd signaling (Figure 7). However, for two genes (*proc* and *FMRFa)*, both of which show strongly enriched neuronal expression, the infection-mediated decreases in mRNA expression were reversed in the *imd* mutants, suggesting that the suppression of these genes may be mediated through Imd signaling (Figure 7). Indeed, Imd signaling has been shown to be induced in neurons through direct effects of peptidoglycan on the brain (Kurz *et al*. 2017).

**Figure 6.**
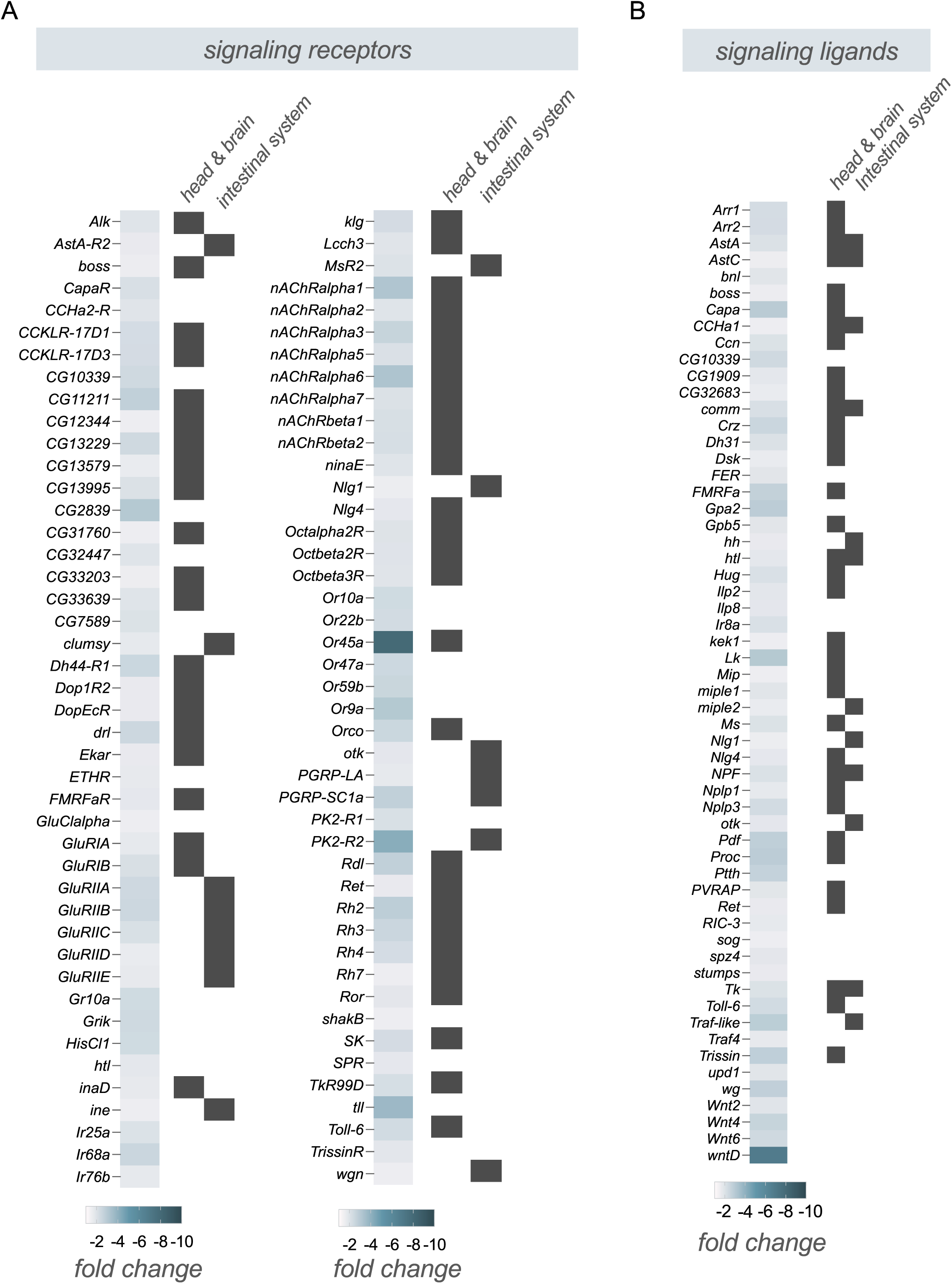
Enteric *P*.*e*. infection downregulates the expression of many signalling receptors and ligands that show brain-and intestine-enriched expression. A, B) Heatmap depicting the change in expression (Log2 fold change vs mock-infected flies) of A, signalling receptors, and B, signalling ligands following enteric infection. Grey symbols show genes with strong enrichment in either the head/brain or intestinal system, based on expression levels from FlyAtlas2.

**Figure 7.**
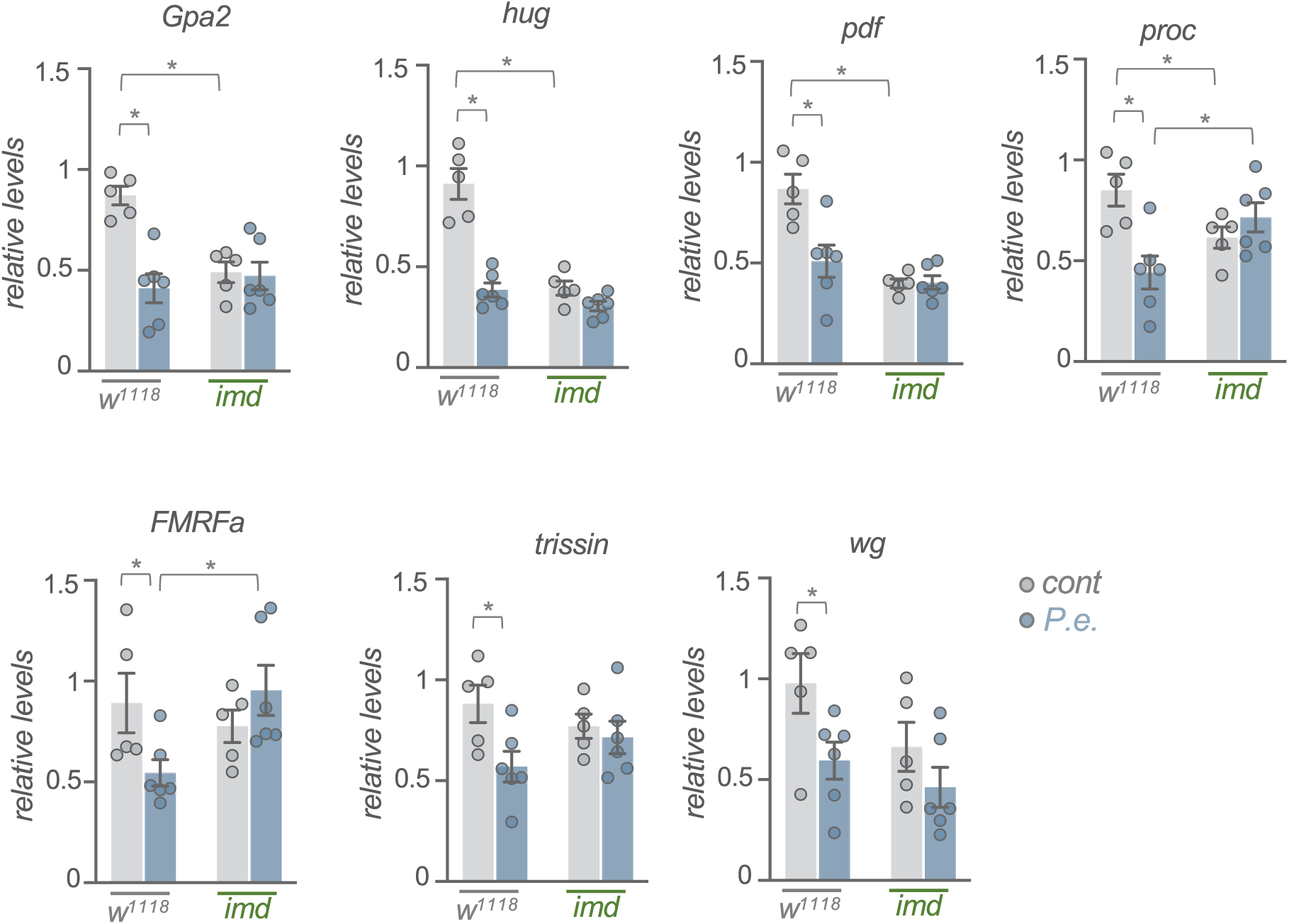
Enteric *P*.*e*. infection downregulates the expression of signalling molecules independently of Imd signalling. qPCR analysis of selected signalling molecule mRNA levels from control (*w*^*1118*^) vs *imd* mutant adult females, that were either mock-infected (grey bars and symbols) or infected by oral feeding with *P*.*e*. for 24 hours (blue bars and symbols). Bars represent mean +/-SEM. Symbols represent individual data points, n=4-6 per condition. * p<0.05, Two-way ANOVA, followed by Students t-test.

Together, both our RNA seq results showing decreased expression of transcription factors and signaling molecules, and the normal tissue expression patterns of these genes from FlyAtlas suggest that enteric infection leads to a) a direct downregulation of both transcription factor and signaling molecule expression within the intestine, and b) a non-autonomous decrease in neuronal signaling. These latter effects may rely on altered gut-to-brain signaling. For example, several of the gut-enriched signaling molecules, such as *AstC, CCHa1, NPF* and *Tk* have receptors in the brain, suggesting they may signal from the gut-to-brain to mediate changes in gene expression. Indeed, a direct gut-to-brain signaling role has been shown for *NPF, AstC* and *Dh31* in the context of nutrient regulation of feeding and metabolism (Yoshinari *et al*. 2021; Kubrak *et al*. 2022; Lin *et al*. 2022). In addition, several of the changes in both transcription factor levels and intestinal/neuronal signaling could explain the increases in metabolic gene expression that we observed. For example, both *sug* and *REPTOR-BP* (both of which we see are downregulated upon infection) have been shown to regulate the expression of both carbohydrate and lipid metabolic genes (Mattila *et al*. 2015; Tiebe *et al*. 2015). Interestingly, *REPTOR-BP*-dependent transcription is negatively regulated by insulin/TOR signaling (Tiebe *et al*. 2015), which would be consistent with some of the changes in metabolic gene expression being mediated though upregulated insulin/PI3K/TOR signaling. Furthermore, several of the intestinal-and neuronally-expressed signaling molecules, such as *NPF* (Yoshinari et al. 2021), *Tk* (Song *et al*. 2014; Kamareddine *et al*. 2018a), *AstC* (Kubrak *et al*. 2022), and *Crz* (Kubrak et al. 2016), have been shown to alter whole-body metabolism (Zhou *et al*. 2020; Medina *et al*. 2022).

One other possibility is that the changes in both intestinal and neuronal signaling molecules that we observed may mediate alterations in fly behaviour upon infection. It is known that infected flies will alter their feeding behaviour, often to avoid eating infected food (Soldano *et al*. 2016; Charroux *et al*. 2020; Cai *et al*. 2021), and they will alter their fecundity and egg-laying behaviour (Masuzzo *et al*. 2019; Masuzzo *et al*. 2020), perhaps to limit the energetically costly process of reproduction while fighting infection (Schwenke *et al*. 2016). Given that several of the neuropeptide signaling molecules and pathways that we see downregulated upon infection are known to affect both feeding and reproduction (Schoofs *et al*. 2017; Ameku *et al*. 2018; Nassel and Zandawala 2019; Hadjieconomou *et al*. 2020; Kim *et al*. 2021; Sadanandappa *et al*. 2021), it is possible that they may modulate either process to help promote infection tolerance. Thus, enteric infection-mediated changes in both gut-to-brain signaling and CNS signaling may be a general mechanism to coordinate multiple whole-body physiological and behavioural responses to infection.

One limitation of our work is that we examined gene expression changes upon infection only in females, and not in males. Interestingly, a recent study using a model of systemic bacterial infection in flies described sex differences in Imd signaling and AMP expression, but not metabolism upon infection (Vincent and Dionne 2021). Thus, it will be interesting to examine whether there are any sex differences in the enteric infection-mediated changes in gene expression and the modulation of these changes by Imd signaling that we see in females.

## Acknowledgements

Stocks obtained from the Bloomington *Drosophila* Stock Center (NIH P40OD018537) were used in this study. We thank Qingrun Zhang for bioinformatic support. This work was supported by CIHR Project Grants (PJT-173517, PJT-152892) and an NSERC discovery grant to S.S.G.

## Author contributions

R.D., and B.L, carried out experiments. R.D., B.L and S.S.G. performed data analysis. S.S.G. directed the study and wrote the manuscript.

## Declaration of interests

The authors declare no competing interests.

### Supplemental Table 1.

List of primers used in this study.

## Methods

### *Drosophila* Stocks and Culturing

The following strains were used: *w*^*1118*^, *imd*^*[EY08573]*^ (Bloomington stock centre #17474). Flies were grown on medium containing 150 g agar, 1600 g cornmeal, 770 g Torula yeast, 675 g sucrose, 2340 g D-glucose, 240 ml acid mixture (propionic acid/phosphoric acid) per 34 L water. All stocks were maintained at either 18°C or RT. For infection experiments, flies were raised from embryos to adults at 25°C. Following eclosion, females were allowed to mate for 2 days before being separated from males and aged for another 5-6 days, at which time infection experiments were performed. All experiments were conducted in mated adult females.

### Enteric infections

Enteric infections were performed using previously described methods (Buchon *et al*. 2010; Zhao and Karpac 2021). Briefly, *Pseudomonas entomophila* (*P*.*e*) from overnight cultures were pelleted and resuspended in 5% sucrose solution (in PBS) such that the final concentration of bacteria was OD_600_= 200. Bacterial pellets were then dissolved in filter sterilized 5% sucrose/PBS. Chromatography paper (Fisher, Pittsburgh, PA) discs were dipped in the bacterial solution and were carefully placed on standard fly food vials such that they covered the entire food surface. Adult females were first subjected to a 2-hr starvation period in empty vials at 29°C. Then 10-12 flies were transferred to each infection vial and then placed in a 29°C incubator for the duration of the 24-hour infection period. Mock-infected control flies were placed in similarly prepared vials that contained paper discs soaked in 5% sucrose/PBS alone.

### Total RNA isolation

Adult flies were snap frozen on dry ice in groups of 5. Total RNA was then isolated using Trizol according to the manufacturer’s instructions (Invitrogen; 15596-018). Extracted RNA was then DNase treated (Ambion; 2238G) to be used for subsequent qPCR or mRNA sequencing.

### mRNA Sequencing

Six independent biological replicates (5 flies per group) of mock infected and *P*.*e*. infected groups were prepared and analyzed. RNA-sequencing was conducted by the University of Calgary Centre for Health Genomics and Informatics. The RNA Integrity Number (RIN) was determined for each RNA sample (6 replicates per each condition were used). Samples with a RIN score higher than 8 were considered good quality, and Poly-A mRNA-seq libraries from such samples were prepared using the Ultra II Directional RNA Library kit (New England BioLabs) according to the manufacturer’s instructions. Libraries were then quantified using the Kapa qPCR Library Quantitation kit (Roche) according to the manufacturer’s directions. Finally, RNA libraries were sequenced for 100 cycles using the NextSeq 500 Sequencing System (Illumina).

### RNA seq analyses

Quality control of sequenced DNA was carried out using FastQC. Sequenced reads were mapped to the *Drosophila* genome Release 6 (GCF_000001215.4_Release_6_plus_ISO1_MT_rna) using Kallisto. Measurements of transcript abundance and differential expression were made using Sleuth.

### Gene Ontology, KEGG pathway, and tissue expression analyses

Analyses of Gene Ontology and KEGG pathway enrichment of up-and down-regulated genes (>1.5-fold, p<0.05) were performed using G-profiler and Revigo. Tissue enrichment of differentially expression in Figures 5 and 6 was examined using FlyAtlas2. We defined genes having enrichment in head and brain (combined expression levels in adult head, eye, brain/CNS, thoracicoabdominal ganglion) and/or intestinal system (combined expression levels in crop, midgut, and hindgut) if expression in these tissues was >3 fold higher than any other tissue(s).

### Quantitative RT-PCR measurements

Total RNA was extracted from either whole flies or from isolated intestines or abdominal samples (abdominal carcass containing attached fat body, but with ovaries, guts and Malpighian tubes removed). The RNA was then DNase treated as describe above and reverse transcribed using Superscript II (Invitrogen; 100004925). The generated cDNA was used as a template to perform qRT–PCRs (ABI 7500 real time PCR system using SyBr Green PCR mix) using gene-specific primers. PCR data were normalized to *5S rRNA* levels. All primer sequences are listed in supplemental table 1.

### Statistical analysis of qRT-PCR data

All qRT-PCR data were analyzed by Students t-test or two-way ANOVA followed by post-hoc students t-test where appropriate. All statistical analysis and data plots were performed using Prism statistical software. Differences were considered significant when p values were less than 0.05.

### Data availability

The RNA-sequence data has been deposited in NCBI’s Gene Expression Omnibus and are accessible through GEO Series accession number GSE202578 (https://www.ncbi.nlm.nih.gov/geo/query/acc.cgi?acc=GSE202578)

---

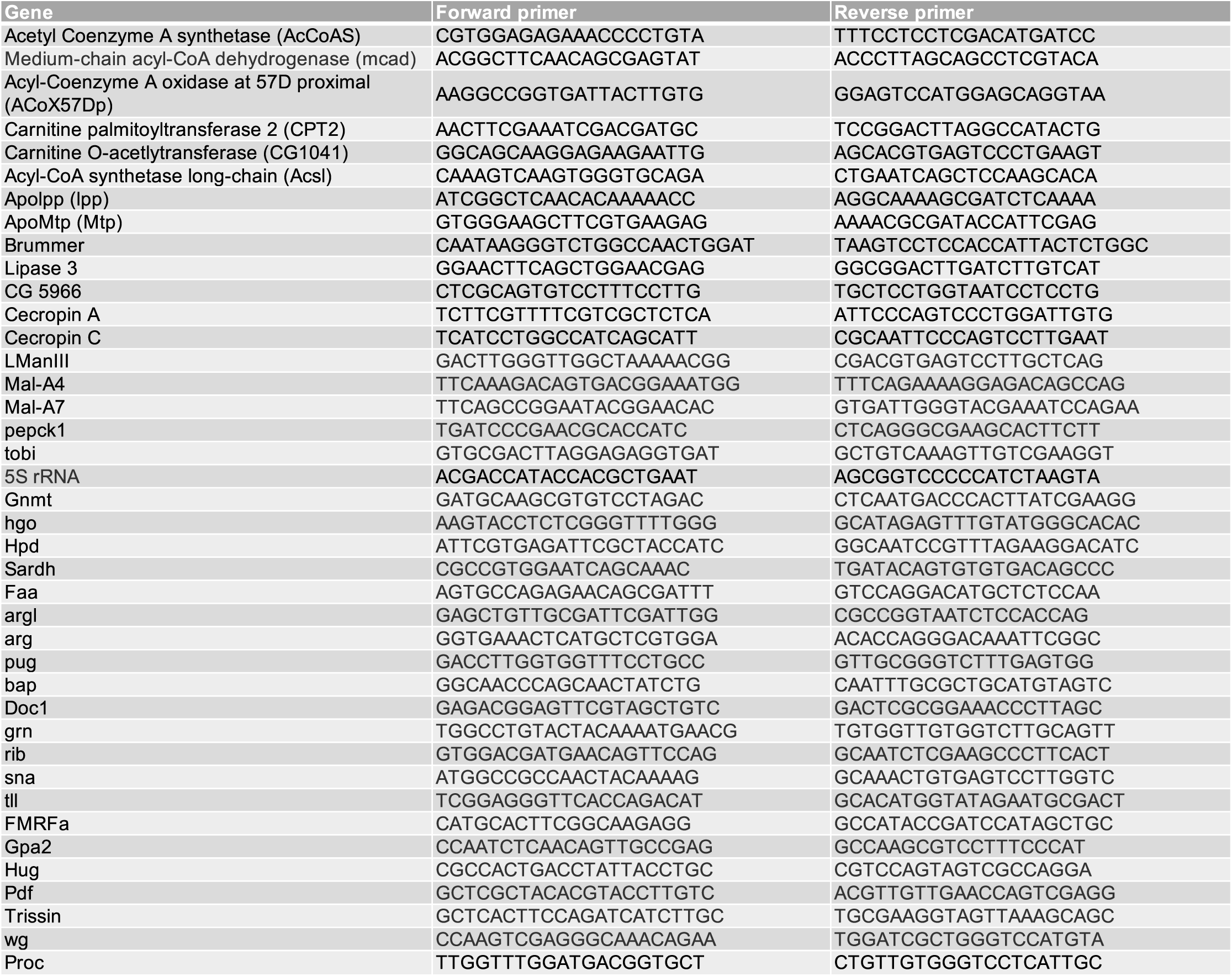

